# SMeta, a binning tool using single-cell sequences to aid in reconstructing species from metagenome accurately

**DOI:** 10.1101/2024.08.25.609542

**Authors:** Yuhao Zhang, Mingyue Cheng, Kang Ning

**Affiliations:** Key Laboratory of Molecular Biophysics of the Ministry of Education, Hubei Key Laboratory of Bioinformatics and Molecular Imaging, Center of Artificial Intelligence Biology, Department of Bioinformatics and Systems Biology, College of Life Science and Technology, Huazhong University of Science and Technology, Wuhan, China

## Abstract

Because of the large volume and complex structure of metagenomic data, traditional binning methods are often hard to classify microbial metagenomes effectively. To deal with these challenges, introducing longer and more accurate single-cell sequencing data is a possible solution. Inspired by the existing MetaBAT2 tool, this study develops a new vector-based binning algorithm, SMeta, which uses both metagenomic and single-cell sequencing data. SMeta is specifically designed for eukaryotic microbial metagenomes, with the long reads characteristic of single-cell data. By introducing the segment tree data structure, the algorithm aligns long single-cell sequences with short metagenomic sequences quickly. This approach allows for the use of reference genomes from genomic databases to replace single-cell data, which makes more precise identification and reconstruction possible for small genome fragments, which are typically overlooked by traditional methods. Also, it might provide a higher purity sequence set for subsequent assemblies.

## Introduction

For environmental and medical microbiology, high-throughput sequencing technologies like next-generation sequencing (NGS) make it possible to get metagenomic genetic information from bacteria, fungi, and other microorganisms within enormous complex samples conveniently. Also, NGS has greatly accelerated the exploration of microbial resources, especially those unknown microorganisms that are difficult to be captured with traditional culturing methods. Also, it gives new ideas for studying microbial ecosystems.

However, the large amount of data also means new challenges. Effectiveness in distinguishing and reconstructing individual microbial full sequences from complex environmental samples has become a challenge for research.

MegaHit[1], EukRep[2], and MetaBAT2[3] are frequently used to process sequence files. MegaHit is a tool designed for rapid and efficient assembly of metagenomic data. By using de Bruijn graph algorithm, it is able to assemble lots of short reads into longer sequences (also called contigs). This tool performs well with sequences with low coverage. It offers advantages including high speed and memory save, and make it a good choice for large-scale metagenomic data assembly. EukRep is used to distinguish between eukaryotic and prokaryotic microbial metagenomic sequences, helping researchers identify the eukaryotic components within complex metagenomic samples. It uses specific algorithms to find unique sequences’ features, and makes the accuracy of eukaryotic genome identification higher than before. MetaBAT2 is also a binning tool for metagenomic contig data, which mainly cluster sequences from mixed samples into several genomes based on sequences’ composition and abundance information like variance and average depth. It utilizes complex algorithms, like conditional probability and distance which is measured by normal distribution, to improve accuracy and efficiency in the binning process.

Although these methods are effective in certain contexts, they are specific for short-read metagenomic data. The large genome sizes and complexities of eukaryotic cells often cause incomplete or inaccurate analysis results. Also, the application of single-cell sequencing offers new ways for assembling eukaryotic sequences. It allows more precise reconstruction of individual cell genomes, and potentially improves binning accuracy. This study is built upon existing algorithms of microbial genome binning tools to develop the SMeta (<u>S</u>egment Tree Based <u>Meta</u>genome Binning Algorithm), specifically optimized for the longer single-cell sequencing reads aligning with metagenomic sequences. We first apply the basic algorithmic framework of tools such as MetaBAT2 and developed a new genome assembly method by using vector comparison and single-cell sequencing FASTA file data. The software is available at https://github.com/YuhaoZhangwow/SMeta.

## Materials and Methodology

### Overview of SMeta

The algorithm takes FASTA files of metagenomic and single-cell sequencing data as input and the binning results for each metagenomic sequence as output.

Tetranucleotide frequency is the frequency of combinations of 4 continuous base pattern in a DNA sequence. Computing tetranucleotide frequencies helps us to understand the 4-kmer characteristics of genomes, and predict different regions in genes, and do comparative genomics work both inside and between species. Considering the complementary nature of DNA, some tetranucleotide patterns can be compressed like AAAA and TTTT, which lower to 136 different tetranucleotide patterns for all DNA sequences. For computational convenience, these frequencies are lined up as a 136-dimensional vector for analysis and comparison of the small differences among genomes. The process is shown in Figure 1.

**Figure 1.**
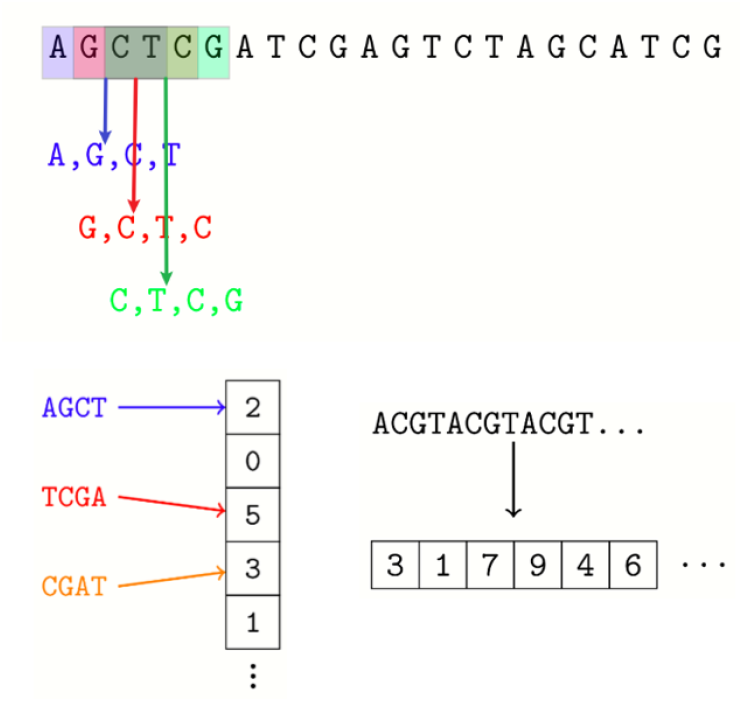
From sequence to vector. Tetranucleotides taken from sliding window on a sequence are 136-class counted and seen as a vector.

The main steps of the SMeta algorithm are as follows: (1) Data reading; (2) Calculating similarity thresholds; (3) Constructing a graph based on the threshold and sequence alignment results; (4) Performing binning using a label propagation algorithm on the graph; (5) Outputting results. The workflow is in Figure 2.

**Figure 2.**
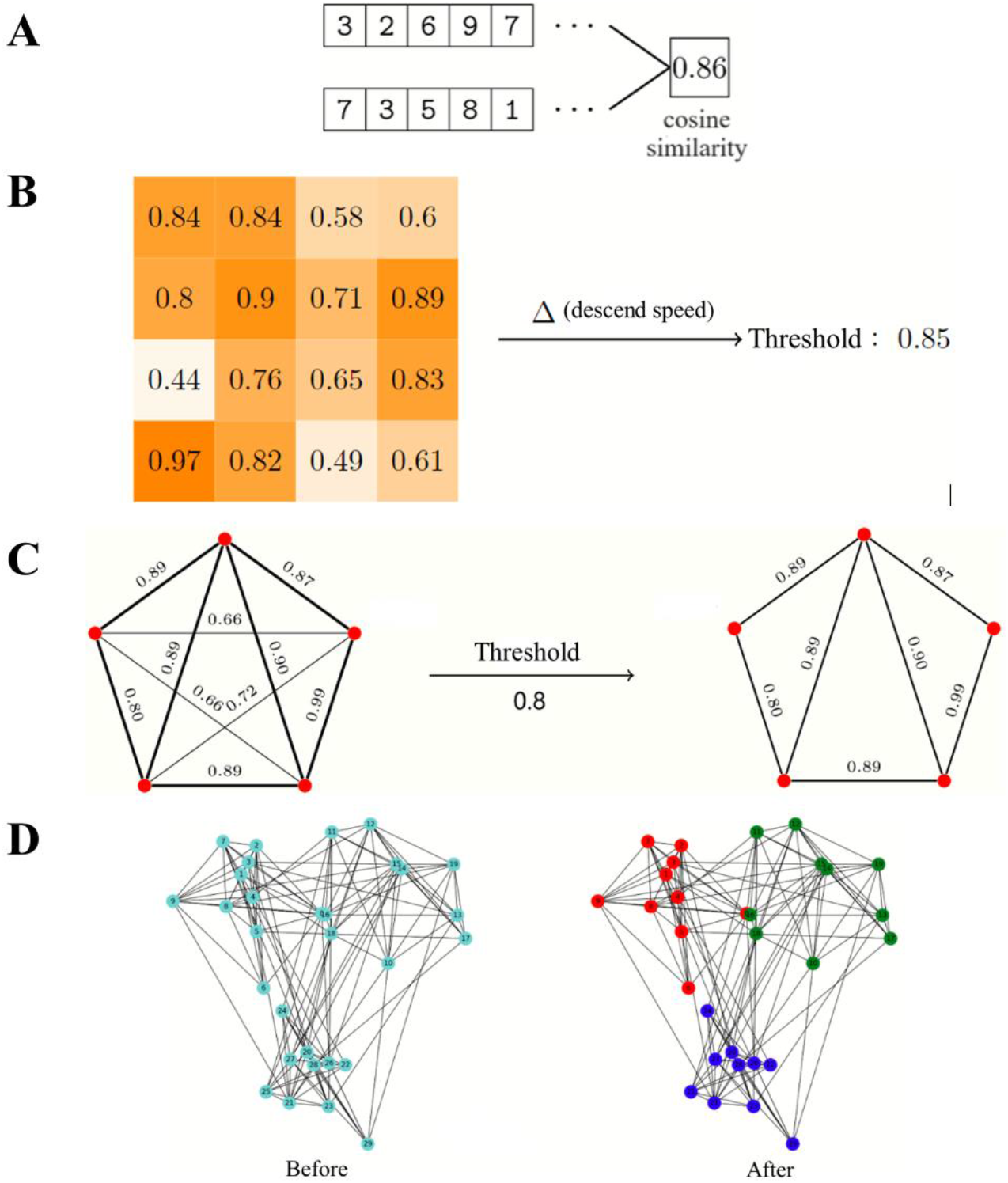
SMeta workflow. (A) Similarity can be quantitatively described by calculating cosine value of included angle of two vectors. (B) Threshold is calculated by similarity matrix, the matrix is the cosine values of all sequence pairs. (C) Graph is built and edges are cut by similarity threshold. (D) LPA is run 10 times for clustering.

**Figure 3.**
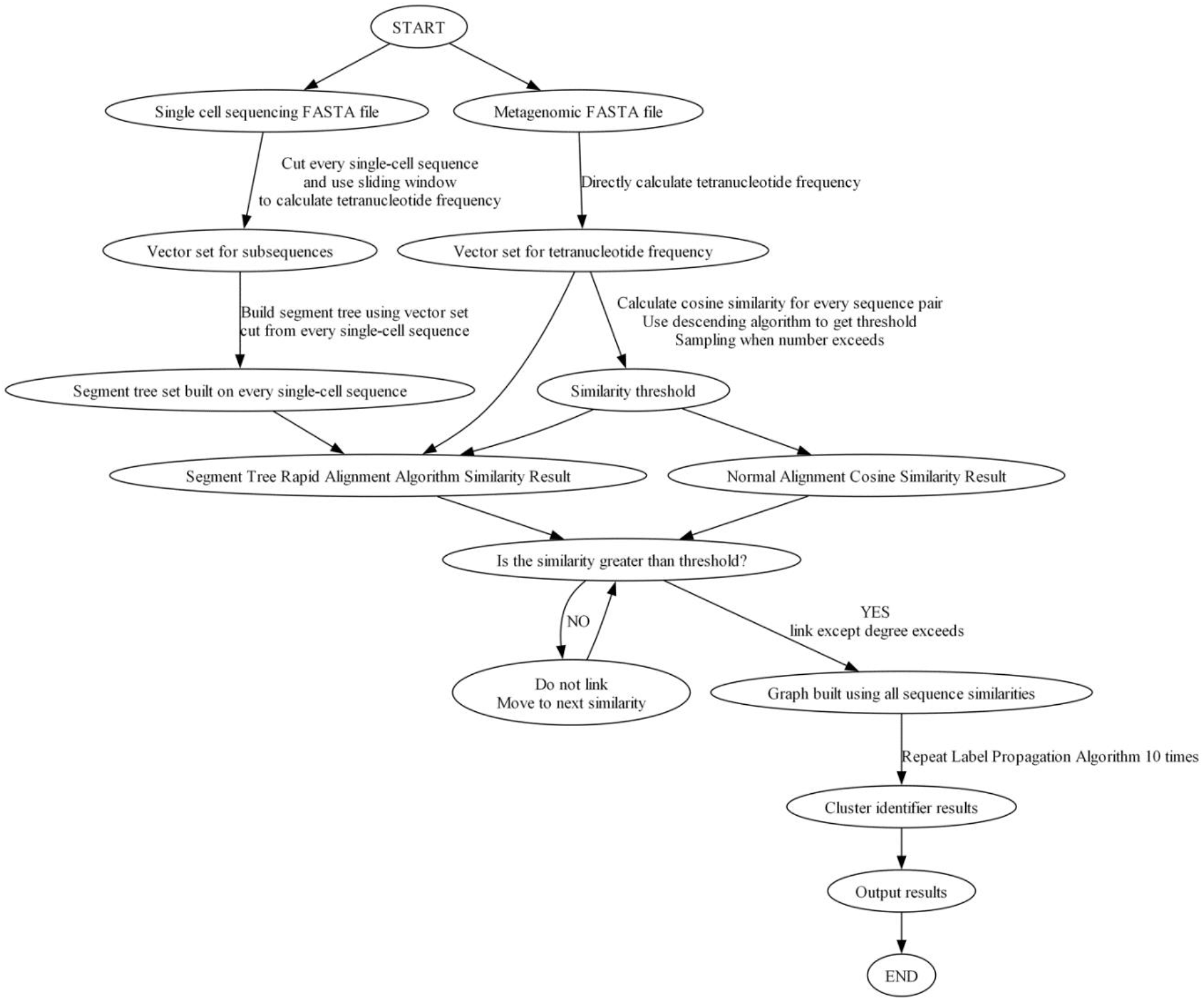
Flowchart of SMeta software: The flowchart describes the pipeline of the software, the input file should follow the standards of FASTA file.

#### 1. Data Reading

In this step, each metagenomic sequence can be turned into a unique tetranucleotide frequency vector. For a long single-cell sequence, direct conversion into just one single vector is a waste of the information it contains. Instead, a sampling rate and sliding window size are used to sample one long single-cell sequence into shorter nucleotide sequences. Each sampled subsequence is turned into a tetranucleotide frequency vector, and it is then used to build a segment tree for rapid alignment with shorter metagenomic sequences. Usually, alignment is done between two sequences, and cosine similarities of the vectors are seen as similarity.

#### 2. Calculating Similarity Thresholds

In this step, the similarity threshold is from using an n×n similarity matrix, which is generated from the comparisons of n metagenomic sequences. A descending method is used to find the similarity threshold for graph construction, which is also demonstrated in the 3^rd^ step with more detail. By setting the threshold, cosine values between sequences below the threshold are not considered, which minimizes the number of edges in the graph and also ensures binning accuracy.

#### 3. Graph Construction Based on Sequence Alignment

This third step is constructing a graph whose edges represent potential binning relationships of sequences. This step can be divided into two parallel processes: constructing the graph based on the similarity between single-cell sequences and metagenomic sequences; constructing the graph based on the internal similarity of metagenomic sequences. The segment tree-based rapid alignment algorithm is used for alignment of long single-cell sequences and short metagenomic sequences. The detail are as follows:

∘ **Internal Similarity of Metagenomic Sequences**: The angles between vectors of each pair of sequences are calculated, making each metagenomic sequence as a vertex in the graph. Edges are added between vertices if their similarity exceeds the threshold. If a vertex has too many edges, only the some (default 200) highest similarity edges are kept, the number is controlled by Maxedge.
∘ **Similarity Between Single-Cell and Metagenomic Sequences**: Each metagenomic sequence and single-cell sequence are seen as vertex. A segment tree is constructed for the sampled subsequences of each single-cell sequence. Each metagenomic sequence A is quickly aligned with the segment tree of each single-cell sequence B. If the alignment results show the similarity is bigger than the threshold, an edge is added between A and B. If a vertex has too many edges, only the top Maxedgeseg (default 50) highest similarity edges are remained.

#### 4. Label Propagation Algorithm for Binning

In the second to last step, the label propagation algorithm[4] is run on the graph to perform binning. Usually, ten times of the label propagation algorithm are run to ensure stability. After the process, each metagenomic sequence is assigned a binning result cluster number.

#### 5. Output Results

In the fifth step, the binning result for each metagenomic sequence is output for further cluster analysis.

### Segment tree rapid alignment algorithm

A segment tree is a binary tree data structure used to save intervals or segments. It divides an interval into many unit intervals, with every indivisible unit interval as a leaf node in the segment tree. The root node represents the entire DNA interval, and the left and right child nodes are the left and right subintervals after equally split the parent node’s interval[5]. A segment tree for an interval set of length n uses O(nlogn) storage space and can be built in O(nlogn) time. It supports searching for intervals of arbitrary length, with a search time complexity of O(logn+k), where k is the number of intervals got. Typically, the average read length of metagenomic sequences is often much smaller than the read length of single-cell sequencing. So a single global alignment between metagenomic and single-cell sequences is meaningless. An obvious approach is using a sliding window on the single-cell sequencing reads, with a window size equal to the length of a metagenomic read, and doing a sequence alignment at each step. However, the classic Smith-Waterman algorithm has a time and space complexity of O(n^2^) for aligning nucleotide sequences of length n. If a single-cell sequencing read is of length m, there will be m−n windows, resulting in a complexity of O(mn^2^), which is unacceptable for single-cell read lengths over 10^5^. Hence, time complexity decrease in alignment and sampling is necessary. Because we need for approximate rather than precise alignment in large data, a segment tree algorithm can lower the complexity of aligning k metagenomic reads with a single-cell sequence to O(m/n+k·log(m/n)).

For sampling, since the precise alignment is unnecessary because the sequences are turned into vectors. The sliding window range can be increased to the average length of the metagenomic reads n. Thus, only ⌈m/n⌉ subsequences are needed, each at positions [1,n],[n+1,2n],… These ⌈m/n⌉ subsequences form a subsequence interval. The average length n is not strictly the mean but the lower quartile of all metagenomic read lengths, which is written as AVGLENGTH. By using the merging properties of segment trees, it allows the algorithm to align a metagenome sequence with several merged intervals. One example is in Figure 4. We here remind again that the alignment is based on tetranucleotide frequency vectors converted from metagenome sequences or subsequence intervals.

**Figure 4.**
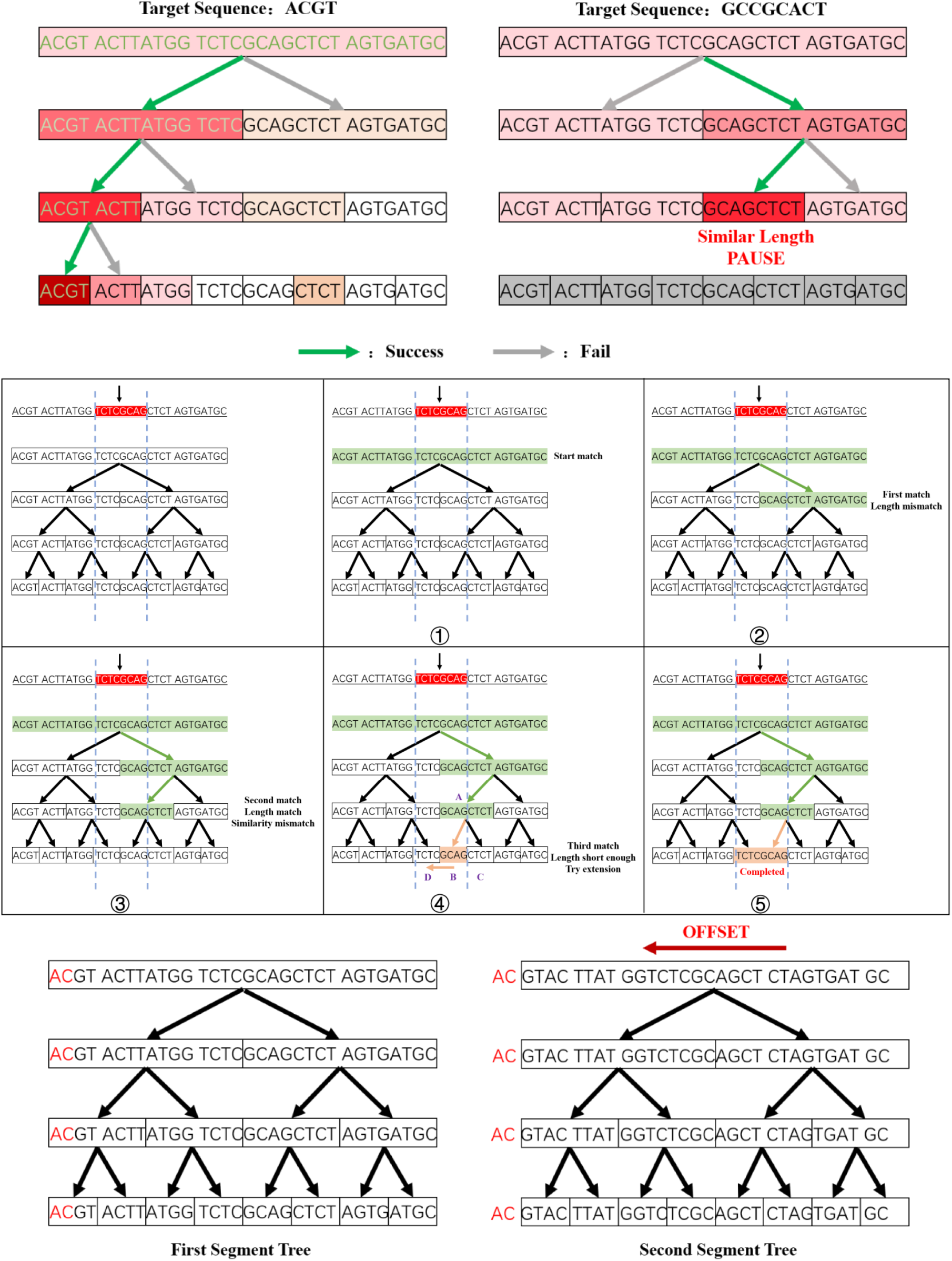
Explanation for Segment tree rapid alignment algorithm. (1) The theoretical assumption for logarithmic complexity. (2) The workflow of one-time extension match operation. (3) The mechanism of offset operation.

The premise of using segment trees is that the intervals must be mergeable, meaning the an interval can be uniquely described by its left and right subintervals. For example, in a segment tree whose intervals describe the sums of the intervals, the parent’s interval’s sum is simply the addition operation of its left and right subintervals’ sums.

For vectors, the merging property is evident: the tetranucleotide frequency vector of a sequence in the interval [l,r] can be calculated by adding the vectors of the intervals [l,mid] and [mid+1,r]. Conversely, L2 normalized vectors cannot do this, as the sum of normalized vectors does not equal the normalized sum vector. Thus, vectors should only be normalized after constructing the segment tree. The merge characteristic of vectors makes quick construction of the segment tree with a time complexity of O(n).

Because the genome sequences remain unchanged, updates within the interval are unnecessary; only query operations are needed. The query operation of the segment tree is to quickly find the most similar single-cell read interval with a given metagenome sequence and ensure the interval length matches the metagenomic read length as much as it can, and then return the highest cosine similarity of the interval vector and metagenome sequence.

A theoretical assumption is that, for a given metagenome sequence, if the interval A in the segment tree is the most similar, and its adjacent interval B is also similar, while the other adjacent intervals C and D are not as similar, then the merged interval AB will be more similar to the metagenomic read than the merged interval CD. Therefore, when the cosine similarity of AB is found to be higher than that of CD, CD and its subintervals can be directly not considered afterwards. This assumption is the theoretical basis for achieving logarithmic complexity in the algorithm. By this approach, the segment tree can deal with interval queries on an interval of length N with a logarithmic time complexity of O(log N), which is much more efficient than linear search O(N) on an array. This fast query capability and the segment tree’s excellent support for interval merging make the segment tree very useful for handling complex interval problems.

For common cases where the expected sequence position of a metagenome sequence across several intervals on a single-cell sequence rather than perfectly aligning with a specific interval, the following extended matching method can be applied if the expected sequence length exceeds twice the AVGLENGTH. When a segment tree interval A of similar length is matched, and the corresponding node of interval A is not a leaf node, while the matching similarity is below the required threshold, an one-time extension match can be attempted. The steps are as follows:

1. Continue matching downwards just once to find the most similar interval in the left and right child intervals B and C of A (assume that B is on the left of C), where the interval length is shorter than the given metagenomic sequence length.
2. If interval B is matched, extend in the opposite direction to B and C to find the adjacent interval D (similarly for C), forming the interval DB. Calculate the similarity between the DB interval and the metagenomic sequence.

However, when dealing with extremely long sequences, it is possible that the most similar subsequence on the single-cell sequence may have a length close to AVGLENGTH and be evenly distributed across two adjacent intervals. In this situation, an interval is split evenly, can result in incorrect calculation of the approximate highest similarity. By simply reducing the sliding step size to solve this issue, it will break the merging characteristic of the intervals in a segment tree, since there would be overlapping parts in the sequences of adjacent intervals, leading to false vector results due to repeated calculations of the overlapping part.

To solve this problem, this method constructs multiple segment trees on a single sequence by offsetting a fixed length, controlled by an optional slice parameter. For example, if there is a single-cell sequence of length m, and the AVGLENGTH value for the metagenome is n, with m being greater than half of n, and the slice parameter is set to 2, then two segment trees need to be constructed. The intervals for the first segment tree would be [1,n], [n+1, 2n], and so on, with no offset. The second segment tree would have an offset of n/2, resulting in intervals such as [0.5n, 1.5n], [1.5n+1, 2.5n], etc. This ensures the merging property of the segment tree is maintained, as the intervals properly align with one another. If the slice parameter is set to another value, 2 can be replaced accordingly.

When the slice parameter is 2, any single-cell subsequence window of length n will be at least 75% covered by the interval of one of the segment trees. That means for a metagenome sequence A, no matter where the exact most similar single-cell sequence interval B is, there will be an interval in one of the segment trees that covers at least 75% of B. This sampling solves the issue of the best interval covering only 50% of a metagenomic sequence.

This offset method significantly reduces the risk of missing during query. During the query, multiple segment trees can be queried in parallel, and the results can be compared to find the most similar subsequence. This enables comprehensive querying and makes the process due to parallel operations quicker.

Additionally, this method of constructing multiple segment trees provides flexibility for subsequent data analysis. For instance, in some cases, the slice parameter size might need to be adjusted because of computational resources limitation and query accuracy. This strategy is particularly suitable for ultra-long and large-scale single-cell sequencing genomic data.

## Results

The software in this paper, SMeta, also supports binning for only metagenomic data. When comparing its performance, we mainly compare it with the classic binning software MetaBAT2. MetaBAT2 is applied to large-scale and complex human microbiome data and can recover hundreds of high-quality genes, including many undetected by other tools. Using coverage files such as bam files, the overall quality can be improved. MetaBAT2 is also highly efficient in computation, because it only needs low memory and shows fast computation speeds. For example, it can perform multiple rounds of binning on large datasets without the need to set the parameters yourself (MetaBAT2 automatically optimizes parameters by default) and can be done with faster speed. For instance, binning a million contigs and approximately 1000 samples spends only a few minutes.

### Data Composition

The data mainly has three types: (1) artificially constructed data of different species to test classification accuracy, (2) metagenomic data, and (3) single-cell sequencing data. The FASTA files of the data primarily focus on metagenomic data, which can be divided into two types:

1. **Artificially Constructed Metagenomic Data:** These are generated by randomly concatenating real single-cell sequencing data, then randomly cutting them into sequences of random lengths. The cuts do not distinguish between different files or sequences, meaning a single sequence may be a combination of several sequences, and noise is added to simulate variations. This data mainly tests the SMeta software’s ability to recover from its own sequences. Figure 5 shows the length distribution of the cut metagenomic sequences, which is a normal distribution with approximately 18,000 sequences.
2. **Real Metagenomic Data from the Tibetan Plateau Water:** The diverse aquatic ecosystems of the Tibetan Plateau, including glaciers, wetlands, hot springs, lakes, and rivers, harbor a rich diversity of aquatic microorganisms, including some uncultivated new lineages[6]. These environments display unique patterns of microbial diversity potential, showing unique adaptations to the extreme conditions of the plateau (e.g., drastic temperature fluctuations, low oxygen concentration, low pressure, and intense ultraviolet radiation). Using metagenomes from such specific conditions helps test the generalization capability of the model. As shown in Figure 5, the majority of sequences in the metagenomic files are short, making comparison speed an important factor when dealing with ultra-long single-cell sequencing sequences.

**Figure 5:**
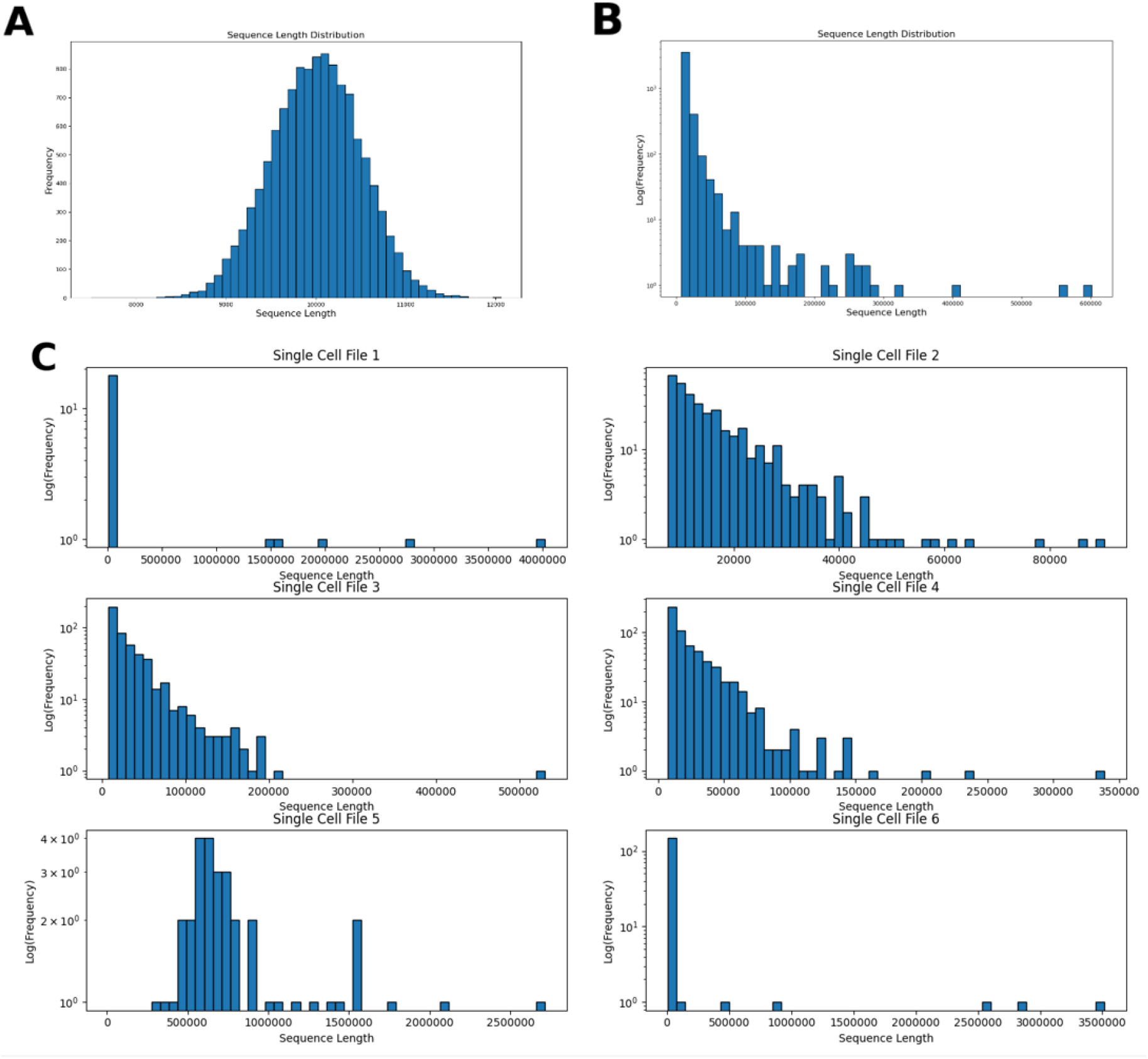
(A) Length distribution of single-cell sequencing data after random cutting and concatenating. (B) Length distribution of metagenomic sequences from the Tibetan Plateau. (C) Length distribution of single-cell sequencing files.

For the single-cell sequencing data from the Tibetan Plateau, six files were selected, with their length distributions shown in Figure 5. Each file contains some long sequences, which is a challenge for SMeta’s fast segment tree alignment algorithm.

### Testing SMeta Algorithm for Correct Classification

Since real datasets do not have classification results to verify the SMeta algorithm, a dataset with known binning results is needed to test the accuracy before applying it to real data. MetaBAT2 is known to be effective; thus, if the SMeta algorithm produces similar results on a dataset with known results, it can be said that the SMeta algorithm can perform basic classification tasks. The dataset consists of five artificially constructed species with a total of 20,000 sequences. To ensure diversity, each species has an average nucleotide mutation rate of 15%, much higher than natural variation rates, with lengths and noise randomly added. The results, shown in Figure 6, indicate that the binning sizes for MetaBAT2 and SMeta across the five species are very close, suggesting that the SMeta algorithm can correctly classify given species, laying the foundation for testing on other datasets.

**Figure 6:**
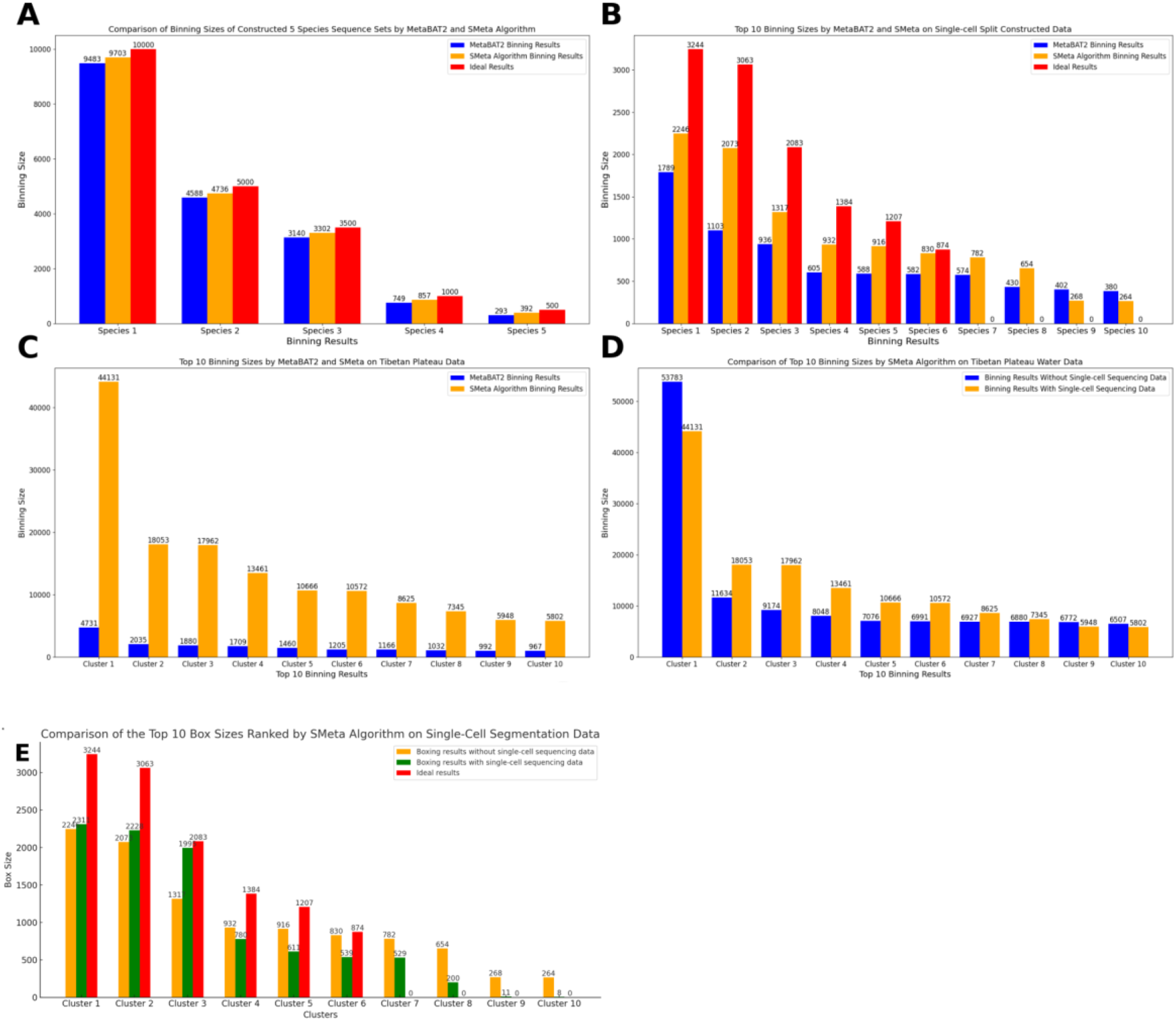
(A) Binning size results of MetaBAT2 and SMeta on constructed five-species dataset. (B) Binning results of MetaBAT2 and SMeta on artificially split and constructed single-cell data. (C) Binning results of benchmark algorithm MetaBAT2 and SMeta on Tibetan Plateau data. (D) Binning results by SMeta before and after introducing single-cell sequencing data on Tibetan Plateau aquatic metagenomic data. (E) Cluster results for comparison before and after introducing single-cell sequences on single-cell split constructed data.

### Comparing Binning Recovery with Other Software Using Metagenomic Data Only

On artificially cut and constructed single-cell data, since the average length of the constructed split sequences is fixed, it is possible to reasonably infer the recovery rate from the lengths of the sequences in different files. After testing, the runtime for both MetaBAT2 and SMeta is almost the same, at 13 seconds. By tracing the representative sequences of the clusters, it is possible to determine the single-cell file to which the cluster belongs. As shown in Figure 6, the recovery performance of SMeta surpasses that of MetaBAT2 for the six single-cell clusters, with recovery rates shown in Table 1.

**Table 1:**
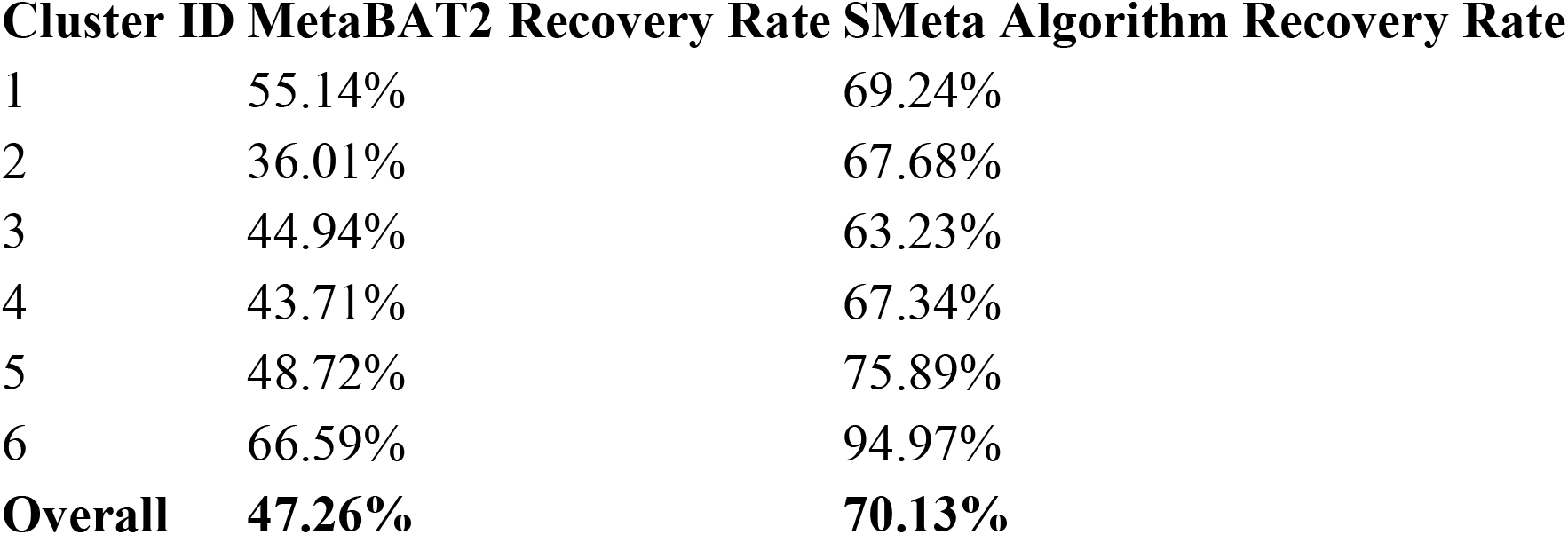
Comparison of recovery rates for the two algorithms.

| Cluster ID | MetaBAT2 Recovery Rate | SMeta Algorithm Recovery Rate |
| --- | --- | --- |
| 1 | 55.14% | 69.24% |
| 2 | 36.01% | 67.68% |
| 3 | 44.94% | 63.23% |
| 4 | 43.71% | 67.34% |
| 5 | 48.72% | 75.89% |
| 6 | 66.59% | 94.97% |
| **Overall** | **47.26%** | **70.13%** |

It is also noted that the bin sizes for SMeta are consistently larger than those of MetaBAT2 for the first eight classifications, indicating that even without single-cell sequencing data, the SMeta algorithm might have better binning capture capability.

For Tibetan plateau data, on the real metagenomic dataset consisting of 184,431 sequences, the SMeta algorithm demonstrates strong detection capability, with the average cluster size being 10 times of that of MetaBAT2. The reason is mainly due to the use of block matrices during the mapping process in the SMeta algorithm, completing the similarity detection in multiple steps, while MetaBAT2 completes it in a single step. By completing it in multiple steps, SMeta allows a sequence to be more thoroughly compared with others, with the maximum number of edges per sequence being about 10 times higher than that of MetaBAT2. MetaBAT2 constructed a total of 586,530 edges, while the SMeta algorithm constructed 64,791,195 edges, making the number of edges in MetaBAT2 less than 1% of that in SMeta. In other words, SMeta uses spatial and temporal costs to achieve extensive connections between sequences, thereby affecting bin sizes. To minimize space and time costs, SMeta maximizes CPU parallelization and block operations to reuse a memory area as much as possible, reducing time and memory overhead.

### Binning Performance of Metagenomic Data with Single-Cell Sequencing Data

For artificially constructed single-cell data, after introducing single-cell sequencing data, the overall recovery rate of the SMeta algorithm reached 71.3%. The most significant result is that sequences from the original clusters 4, 5, and 6 were reclassified into the top three clusters, with cluster 3 seeing the most increase, possibly indicating some similarity among clusters 3, 4, 5, 6. This also suggests that the introduction of single-cell sequencing data influences the labels. Also, the first six clusters were larger, with the significant reduction in the sizes of other clusters, and the sizes of clusters from the ninth can be nearly overlooked.

For Tibetan plateau data, After introducing single-cell sequencing data, the most significant result was that the sequences in cluster 1 increased by 21.9%, while clusters 2-8 saw a decline. This suggests that sequences in cluster 1 might have high similarity to a single-cell file, providing more sequence references for subsequent assembly operations. The decrease in other clusters indicates that sequences in these clusters might also have some similarity to cluster 1, suggesting possible similar species, requiring additional phylogenetic analysis.

In the extreme stress test of sequence alignment between single-cell sequences and metagenomic sequences, using the segment tree fast alignment algorithm, the speed reached 1,350,000 alignments per second, showcasing the superiority of the SMeta algorithm.

## Discussions

Based on the analysis presented, we can conclude that under the premise of accurate classification, the SMeta tool is better than MetaBAT2 in pure metagenomic binning tasks. Additionally, the introduction of single-cell sequencing data results in more clustered bins, and the comparison before and after binning suggests similarities among species. This paper mainly explores the design of the SMeta algorithm, focusing on assembling metagenomic and single-cell sequences. Improving binning effectiveness is the key issue addressed by the SMeta algorithm, leading to the core problem of aligning long and short sequences in metagenomic and single-cell sequence comparisons. This study mainly achieved the following algorithm designs:

1. Utilizing vectors corresponding to tetranucleotide frequencies and using the angle between vectors to represent similarity significantly reduces computation and facilitates the parallel execution of the SMeta program.
2. Observing the additive property of vectors and combining it with the linear structure of sequences led to the application of segment tree data structures to solve related issues.
3. Employing binary search reduces the search space, and establishing multiple segment trees on a single sequence by using sequence offsets increases sequence coverage, achieving more comprehensive alignments with minimal spatial cost. While the SMeta algorithm presents certain improvements over the classical MetaBAT2 in terms of design and implementation, there remain many challenges and areas for improvement in the system and code. This study identifies the following areas for future enhancement:
4. The dataset scale used in this study is limited; running SMeta on files larger than 1GB (approximately 500,000 metagenomic sequences) often leads to memory insufficiency, causing the program to crash after several hours.
5. The SMeta algorithm operates without prior information over the sequences themselves, making it incapable of identifying enhancers, promoters, and translocation events. These can only be inferred retrospectively based on tetranucleotide frequency relationships.
6. The components of the tetranucleotide frequency vector are mutually orthogonal and independent, lacking the ability to capture variations, especially in cases of large-scale inversions, insertions, and deletions.

In conclusion, this study designs the new SMeta algorithm, integrating metagenomic and single-cell data to improve binning operations and provide better sequence data for hybrid assembly. Tests on different datasets demonstrate the algorithm’s effectiveness in enhancing binning performance and assembly accuracy. The introduction of single-cell data particularly strengthens homology recognition among sequences, which is crucial for understanding complex microbial communities. Future work will focus on optimizing the technical challenges of the SMeta algorithm and software while exploring the use of biological characteristics to further enhance the algorithm’s recognition capability. This SMeta algorithm research offers new perspectives and tools for metagenomic and single-cell sequence analysis, showcasing the potential applications of computational biology in modern biological research.

## Acknowledgements

Numerical computations were performed on the Hefei Advanced Computing Center.

## Contributors

YHZ designed the algorithm, analyzed the data, completed software development, debugged, and drafted the manuscript. KN and MYC designed the study, collected clinical data, and interpreted the data, drafted and edited the manuscript, and supervised the study. All authors critically reviewed the manuscript.

